# Deciphering Parasitic Strategies: Dual Transcriptomics Reveal Distinct Infection Mechanisms and Gall-like Traits in Rafflesiaceae

**DOI:** 10.64898/2026.06.28.735044

**Authors:** Marco Bürger, Adhityo Wicaksono, Susan Pell, Allen Mamerto, Todd P. Michael, Jeanmaire Molina

## Abstract

Rafflesiaceae, known for producing the largest flowers in the world, are obligate parasites that exclusively infect *Tetrastigma* sp. (Vitaceae). Despite their unique biology, the interactions between parasitic tissues and host roots remain poorly understood, particularly during the flower morphogenesis phase, where parasitic tissue erupts through the host root. Here, we performed dual transcriptome analyses of two Rafflesiaceae species and their respective *Tetrastigma* hosts: *Sapria himalayana* with *T. cauliflorum* and *Rafflesia speciosa* with *T. magnum*. Our findings reveal species-specific transcriptional responses in *Tetrastigma,* suggesting divergent parasitism strategies between *Rafflesia* and *Sapria*. Moreover, we identify molecular signatures of parasitism that parallel plant gall formation, particularly in genes governing cell wall modification and host tissue reorganization. Unlike bacterial or insect-induced galls, these mechanisms may involve fungal symbionts, highlighting the unique nature of these interactions. Together, our results demonstrate that Rafflesiaceae parasitism represents a complex tripartite relationship among host, holoparasite, and associated microbes, offering new insights into the hidden biology of these remarkable parasitic plants.

## Introduction

Rafflesiaceae, known for producing the world’s largest flowers, are a parasitic plant family that exclusively parasitizes *Tetrastigma* (Vitaceae). As endophytic holoparasites, Rafflesiaceae lack stems, roots, and leaves, and subsist entirely inside their host for nutrient acquisition until flowering. The parasitic tissue infects the host root at the vascular bundles and resides within the vascular cambium and xylary regions [1,2]. These tissues proliferate as clusters of cells or uniseriate strands [1,3], which can remain dormant until an unknown trigger [4] induces prolific cell division, culminating in the eruption of a flower bud. During this process, significant alterations occur in the host’s vascular tissue as the flower bud enlarges, including bending and compression of the xylem [5]. Despite these observations, the molecular processes and cellular impacts involved in Rafflesiaceae-host interactions remain unexplored.

Several studies highlight the advancements in understanding parasitic plant-host interactions through transcriptomic analyses, revealing key parasitism-related genes and molecular strategies. Parasitic plants such as *Cuscuta* and Orobanchaceae exhibit genes involved in cell wall modification, hormone biosynthesis, and immune suppression, which are central to their parasitic lifestyle [6–8]. For example, *Cuscuta* secretes enzymes like pectin methylesterase (PME) to infiltrate host tissues, while Orobanchaceae parasites express “core parasitism genes” coopted from root and floral tissues in their non-parasitic ancestors [8]. Transcriptomic approaches have also revealed unique gene expression patterns in parasitic plants, distinguishing them from non-parasitic relatives and uncovering adaptations for parasitism [7].

Expanding on these insights, integrated metabolomic and transcriptomic approaches identified key parasitism strategies in *Cuscuta japonica*, uncovering distinct interactions with host and non-host plants [9]. Further studies revealed regulatory modules driving host cell wall degradation during *C. campestris* invasion [10] and the role of host-produced ethylene in facilitating cell expansion and endoreduplication in dodder search hyphae [11]. Additionally, interspecific signaling mechanisms regulating xylem vessel differentiation in *C. campestris* haustoria were discovered [12]. Collectively, these studies underscore the power of transcriptomic analyses in uncovering the molecular mechanisms of parasitism, paving the way for applying dual RNA-seq to holoparasitic systems like Rafflesiaceae.

Recent transcriptomic studies of gall systems have revealed striking parallels with plant parasite-host interactions, offering valuable insights into the molecular mechanisms underlying host manipulation. Gall-inducing organisms, such as insects and fungi, reprogram host plant gene expression to create specialized structures that support their development—much like parasitic plants manipulate their hosts for nutrient acquisition and growth. Both gall formation and parasitic invasion involve hormonal reprogramming, characterized by the upregulation of auxins, cytokinins, and gibberellins that promote cell proliferation and differentiation [13,14]. In *Rafflesia*, the presence of gallic acid derivatives, gall-associated bacterial families, and adenine – a cytokinin linked to gall formation – further supports the hypothesis that *Rafflesia* buds operate similarly to plant galls, manipulating host tissues to facilitate reproductive development [15]. Genes associated with cell wall remodeling, such as those encoding pectin methylesterase and expansins, are also upregulated in both systems, facilitating host tissue infiltration and modification [10,16]. Additionally, metabolic reprogramming occurs as host resources are redirected to support the gall or parasite, and host defense mechanisms are suppressed to ensure successful establishment [6,9]. The activation of reproductive and developmental programs in galls mirrors the strategies of parasitic plants like *Cuscuta* and Rafflesiaceae, which co-opt host tissues to develop their own structures [1,17]. These similarities suggest that parasitic plants such as Rafflesiaceae may employ analogous molecular strategies to manipulate their hosts, highlighting the potential of dual transcriptomic studies to uncover the mechanisms of their unique parasitic lifestyle.

Dual transcriptomics is a powerful tool for simultaneously capturing gene expression profiles of interacting organisms, providing critical insights into host-pathogen dynamics [18]. While extensively applied to human-pathogen studies and some plant systems, its application to parasitic plant-host interactions, particularly holoparasitic systems, remains largely unexplored. Investigating the dual transcriptome of Rafflesiaceae and their hosts presents an unprecedented opportunity to uncover the molecular mechanisms underlying parasitism in one of the most enigmatic plant families. While vascular deformation and tissue remodeling in *Tetrastigma* roots during *Rafflesia* bud development suggest host manipulation, the genes and pathways involved are uncharacterized [5]. Furthermore, the lack of visible stress on host tissues during the vegetative stage of the endophyte underscores the need to explore the molecular strategies enabling parasite establishment without triggering strong host defenses. Studies of *Cuscuta* and Orobanchaceae have revealed key immune-related genes, such as SERK1, SOBIR1, and HaOr7 [19], which are central to host-parasite interactions in these lineages and may similarly play a role in Rafflesiaceae-host interactions. Despite the potential of dual transcriptomics, its application to Rafflesiaceae systems is hindered by limited genomic resources. To date, the genomes of *Sapria himalayana*, *Tetrastigma voinierianum,* as well as *T. hemsleyanum* have been published [20,21], however, either assemblies or annotations are unavailable. Expanding genomic annotations and applying dual RNA-seq to this system could provide insights into the molecular underpinnings of parasitism, including haustorium formation, nutrient acquisition, and host manipulation. This study aims to address these gaps by employing dual transcriptomics to analyze gene expression in Rafflesiaceae and their hosts at the host-bud interface. By identifying the molecular pathways driving host manipulation and parasitic adaptation, this research advances our understanding of Rafflesiaceae biology and the broader evolution of parasitism in plants.

## Results

### De novo genome assembly of *Tetrastigma magnum*

To generate a reference genome for mapping of *Tetrastigma* transcripts, we sequenced the genome of *Tetrastigma magnum*. Two separate sequencing runs yielded a combined total of 217.81 Gb of HiFi reads with a median read quality of Q34. The HiFi reads had a mean length of 16,797 bp and a median length of 15,822 bp (averaged from the two runs). From the sequencing runs, 13,494,375 ZMWs passed filters (35.12% of input), resulting in 12,967,644 HiFi reads. Of the total bases sequenced, 91.5% (199.23 Gb) had quality scores ≥Q30. The coverage depth was 48x. The final assembly size was 2.38 Gb, consisting of 317 contigs with a contig N50 of 75.27 Mb. A total of 47,190 coding sequences (CDS) were identified, and BUSCO analysis revealed 94.4% complete BUSCOs (Supplementary Table 1).

### RNA-seq mapping to *Tetrastigma* and *Sapria* reference genomes

For RNA-seq mapping to the *Tetrastigma magnum* reference genome, uninfected *T. magnum* (total reads: 191,919,296) yielded 68.85% (132,136,935 reads) uniquely mapped reads with 9.36% (17,963,646 reads) multi-mapping reads and a mismatch rate of 0.42%, while *Rafflesia*-infected *T. magnum* (total reads: 376,724,846) showed 42.81% (161,276,507 reads) uniquely mapped with 6.72% (25,316,310 reads) multi-mapping reads and 0.36% mismatch rate. Cross-species mapping resulted in uninfected *T. cauliflorum* (total reads: 181,320,714) showing 33.96% (61,576,515 reads) uniquely mapped and 3.55% (6,436,885 reads) multi-mapping reads with a 3.03% mismatch rate, while *Sapria*-infected *T. obovatum* (total reads: 334,651,464) resulted in 26.76% (89,552,732 reads) uniquely mapped and 3.12% (10,441,126 reads) multi-mapping reads with a 2.23% mismatch rate. When mapping to the *Sapria himalayana* reference genome, the *Sapria*-infected *T. obovatum* sample showed 38.55% (129,008,139 reads) uniquely mapped, 3.08% (10,307,265 reads) multi-mapping reads with a 0.41% mismatch rate, while the *Rafflesia*-infected *T. magnum* sample yielded 31.45% (118,479,964 reads) uniquely mapped and 10.13% (38,162,927 reads) multi-mapping reads with a 3.12% mismatch rate (Supplementary Table 2).

### Transcriptional responses in *Tetrastigma* contrast between *Rafflesia* and *Sapria* infections

We analyzed the number of differentially expressed genes in *Tetrastigma* upon infection by *Rafflesia* vs *Sapria* and found substantial overlap, with 7,021 shared genes, while *Rafflesia*-infected and *Sapria*-infected *Tetrastigma* samples showed 7,990 and 3,784 unique differentially expressed genes respectively (Figure 1A). We then examined the top 20 differentially expressed genes in *Tetrastigma* hosts to gain insight into possible distinct transcriptional responses to infection by *Rafflesia* and *Sapria*. In *Rafflesia*-infected tissues, we observed a strong trend towards upregulation, with 17 out of the top 20 genes showing increased expression and 3 genes being downregulated (Figure 1B). The most highly upregulated gene (log2FoldChange of 12.23) encoded a BURP domain protein (IPR004873). Other significantly upregulated genes included those encoding zinc finger BED-type proteins (IPR003656), alpha/beta hydrolase fold proteins (IPR013094), glycoside hydrolases (IPR001764), and sugar transporters (IPR003663). Among the downregulated genes in *Rafflesia* infection were several defense-related genes, including cystatin domain proteins (IPR000010) and Toll/interleukin-1 receptor homology (TIR) domain-containing proteins (IPR000157), with log2FoldChanges ranging from -8.67 to -9.43. In *Sapria* infection, we observed an even stronger trend towards upregulation, with 19 out of the top 20 differentially expressed genes showing increased expression and only 1 gene being downregulated (Figure 1C). The most highly upregulated gene (log2FoldChange of 10.19) encoded an Early nodulin 93 ENOD93 protein (IPR005050). Other significantly upregulated genes included those encoding glutathione S-transferases (IPR004045), NADH:ubiquinone oxidoreductases (IPR001457), and cytochrome c oxidase subunits (IPR000298). The single downregulated gene in the top 20 for *Sapria* infection encoded a pentatricopeptide repeat protein (IPR002885) with a log2FoldChange of -8.23.

**Figure 1:**
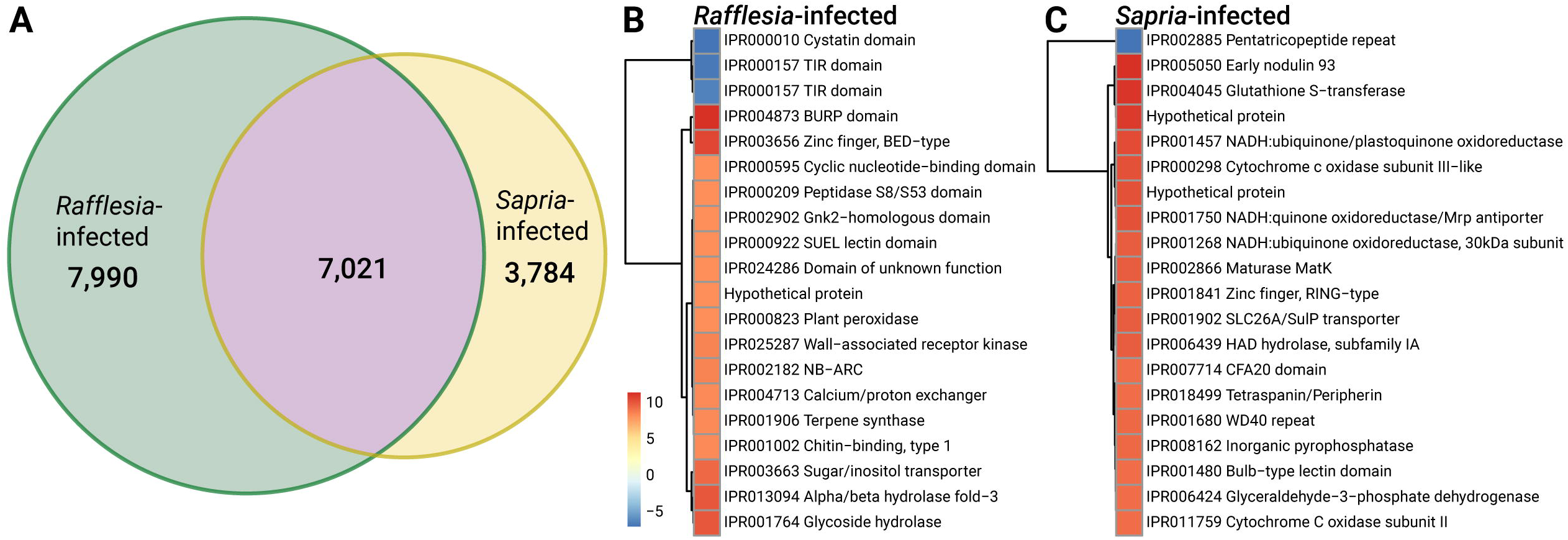
Differential gene expression in *Tetrastigma* response to *Rafflesia* and *Sapria* infection. **A:** Euler diagram showing overlap of differentially expressed genes between *Rafflesia*-infected (green) and *Sapria*-infected (yellow) samples. Numbers indicate unique and shared differentially expressed genes. **B,C:** Heatmaps showing log2 fold changes of top 20 differentially expressed genes in *Rafflesia*-infected (B) and *Sapria*-infected (C) *Tetrastigma* samples. Red indicates upregulation and blue indicates downregulation. Genes are labeled with their InterPro domains and protein names, and clustered by expression pattern.

We conducted COG analysis to characterize shared and distinct patterns of gene regulation in *Tetrastigma* in response to *Rafflesia* and *Sapria* infections (Figure 2A). In *Rafflesia* infection, nearly all COG categories showed strong upregulation. The categories with the highest number of upregulated genes included function unknown (S; 3,751 up vs. 293 down), signal transduction mechanisms (T; 1,224 up vs. 176 down), and posttranslational modification, protein turnover, chaperones (O; 942 up vs. 128 down). Other categories showing substantial upregulation included transcription (K; 907 up vs. 106 down), translation, ribosomal structure, and biogenesis (J; 684 up vs. 20 down), and carbohydrate transport and metabolism (G; 653 up vs. 52 down). The downregulated response in *Rafflesia* infection was relatively modest, with the highest number of downregulated genes found in function unknown (S; 293 down), signal transduction mechanisms (T; 176 down), and posttranslational modification, protein turnover, chaperones (O; 128 down). In *Sapria* infection, while upregulation still predominated, the response was more moderate in magnitude with a higher proportion of downregulated genes. The categories with the highest number of upregulated genes were function unknown (S; 1,525 up vs. 1,337 down), posttranslational modification, protein turnover, chaperones (O; 611 up vs. 263 down), and transcription (K; 453 up vs. 323 down). Several categories showed nearly balanced up- and downregulation, such as signal transduction mechanisms (T; 519 up vs. 512 down). The infection also triggered stronger downregulation in some categories, notably secondary metabolites biosynthesis, transport, and catabolism (Q; 151 up vs. 168 down) and defense mechanisms (V; 44 up vs. 53 down). In summary, while both parasitic plants induced widespread transcriptional changes, *Rafflesia* infection generally elicited a stronger upregulatory response compared to the more balanced regulation pattern seen in *Sapria* infection.

**Figure 2:**
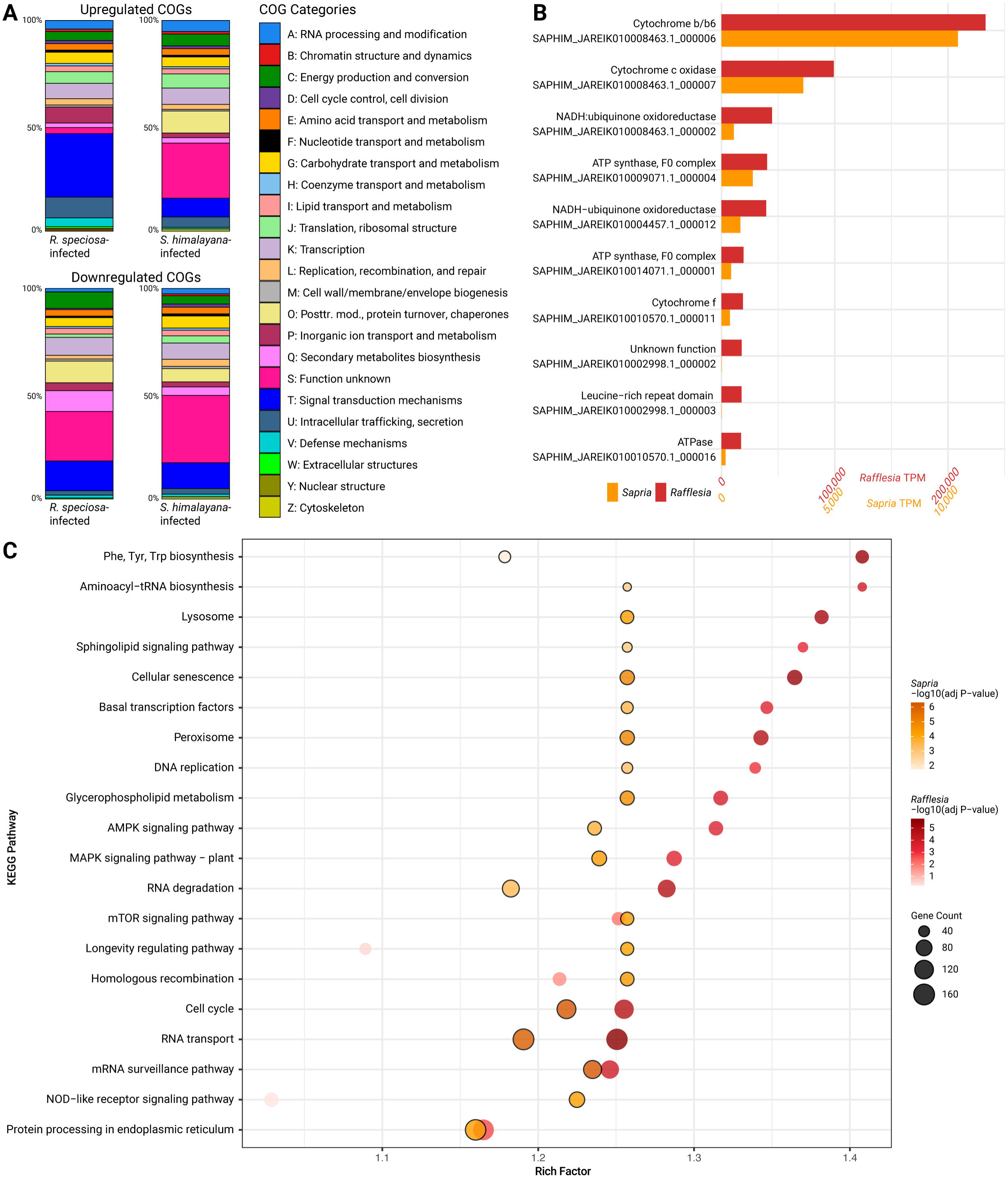
Differential gene expression and functional enrichment analysis. **A:** Distribution of differentially expressed genes by Clusters of Orthologous Groups (COG) functional categories in *R. speciosa*-infected and *S. himalayana*-infected *Tetrastigma* samples. Stacked vertical bars show upregulated (top) and downregulated (bottom) genes, with colored segments indicating the number of genes per COG category. **B:** Bar plots showing the expression levels of the top 10 most highly expressed genes in *Rafflesia speciosa* (red bars) and *Sapria himalayana* (orange bars), measured in transcripts per million (TPM). Each bar corresponds to a gene identified by its function and gene ID. **C:** KEGG pathway enrichment analysis showing significantly enriched pathways for differentially expressed genes in *Rafflesia* and *Sapria*. Dot size represents gene count, color intensity indicates statistical significance (-log10 adjusted *p*-value), and rich factor represents the ratio of differentially expressed genes to total genes in each pathway. Orange dots represent *S. himalayana*-infected samples and red dots represent *R. speciosa*-infected samples.

We subsequently performed GO enrichment analysis to investigate distinct patterns between *Sapria* and *Rafflesia* infection responses in *Tetrastigma*. *Sapria* infection triggered extensive enrichment of GO terms, particularly those involved in vesicle-mediated transport, protein catabolism, and ubiquitin-dependent processes. The most enriched cellular components included various membrane-associated structures and vesicles, suggesting active membrane trafficking during *Sapria* infection. In contrast, *Rafflesia* infection showed limited GO term enrichment, with only cytosolic ribosome being significantly enriched, suggesting a more subtle impact on host cellular processes (Supplementary Figure 1).

### *Rafflesia* and *Sapria* Exhibit Distinct Gene Expression Profiles and Pathway Enrichments During Host Infection

We investigated gene expression differences in the parasitic plants by analyzing the top 10 highly expressed genes that are shared between *Rafflesia* and *Sapria* during host infection (Figure 2B). Both parasites showed consistent expression of genes involved in energy production and metabolic processes, though with notable differences in expression levels. The most highly expressed shared gene in both species was cytochrome b/b6, though *Rafflesia* showed higher expression levels (TPM: 233,124) compared to *Sapria* (TPM: 10,436). Similarly, cytochrome c oxidase subunit I was the second highly expressed gene in both species, with *Rafflesia* again showing much higher expression (TPM: 99,157 vs. 3,611 in *Sapria*). Other consistently shared genes included NADH:ubiquinone oxidoreductase subunits and ATP synthase complex components, all showing the same pattern of higher expression in *Rafflesia* compared to *Sapria*. Both species also expressed genes with leucine-rich repeat domains, which are often associated with plant defense responses, though again at much higher levels in *Rafflesia* (TPM: 17,737) than in *Sapria* (TPM: 7). The shared expression of genes involved in energy production suggests a fundamental importance of energy metabolism during host infection in both parasites, while the consistently higher expression levels in *Rafflesia* may reflect differences in metabolic demands or infection strategies between the two species.

We then analyzed KEGG pathway enrichments to compare patterns between *Rafflesia* and *Sapria* during host infection (Figure 2C). In *Rafflesia*, the most enriched pathways included phenylalanine, tyrosine, and tryptophan biosynthesis, aminoacyl-tRNA biosynthesis, and glutathione metabolism, which were primarily related to amino acid synthesis and antioxidant defense. Other highly enriched pathways such as the sphingolipid signaling pathway, and the citrate cycle (TCA cycle) indicated a significant involvement in energy production and lipid metabolism [22]. In *Sapria*, the top enriched pathways were glycerophospholipid metabolism, peroxisome, cellular senescence, and the mTOR signaling pathway, which were associated with membrane biogenesis, oxidative stress response, and cell growth regulation [23]. Additionally, the enrichment of the NOD-like receptor signaling pathway suggests an involvement in immune response mechanisms [24]. Both parasites showed enrichment in RNA-associated processes, including mRNA surveillance, RNA transport, and RNA degradation pathways, suggesting active regulation of RNA metabolism during host infection.

### Pathway-specific host gene expression during *Rafflesia* and *Sapria* infection

We analyzed gene expression patterns associated with various biological pathways in *Tetrastigma* during infection by *Rafflesia* and *Sapria*. In terms of cell wall integrity, *THE1, FER* and *MCA1* showed strong upregulation during *Sapria* infection, while *WAK* was notably downregulated (Figure 3A). For plant-plant interaction genes, *Or7* was significantly upregulated in both infection contexts, with a higher fold change in *Rafflesia* infection. *SOBIR1* showed strong upregulation in *Sapria* infection but was relatively unchanged in *Rafflesia* (Figure 3B). Among plasmodesmata-associated genes, *CALS1* was upregulated in both infections, with a higher fold change during *Sapria* infection, while *BAM* showed upregulation in *Sapria* but downregulation in *Rafflesia* (Figure 3C). Within the strigolactone biosynthesis pathway, most genes were downregulated in both infections, with *DWARF27* showing the most pronounced downregulation in *Sapria*. Notably, *MAX1* was downregulated in *Rafflesia* but slightly upregulated in *Sapria* (Figure 3D). For xylem formation, *KNAT7* and *SND1* were upregulated in *Sapria* but not in *Rafflesia*, while *VND6* and *VND7* were strongly downregulated in *Sapria* but upregulated in *Rafflesia* (Figure 3E). These findings highlight distinct transcriptional responses in *Tetrastigma* depending on the infecting parasitic plant, suggesting different infection strategies or host responses between *Rafflesia* and *Sapria*.

**Figure 3:**
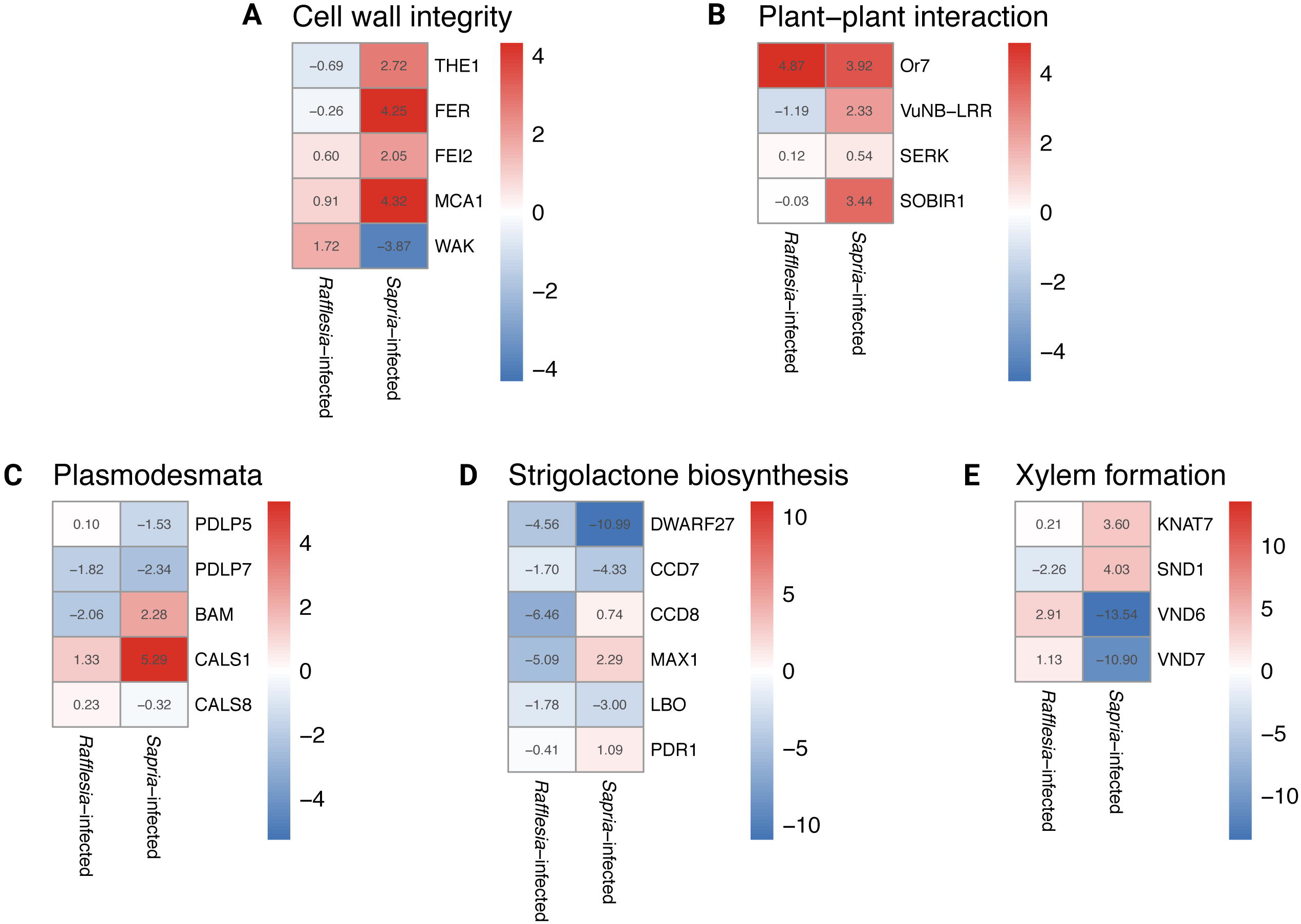
Heatmap comparison of groups of functional genes differentially expressed in *Tetrastigma* infected by *Rafflesia* vs *Sapria*. The heatmap displays log2 fold changes in gene expression, with red indicating upregulation and blue indicating downregulation. Each row represents a gene. The figure includes heatmaps for five functional groups: Cell wall integrity, plant-plant interaction, plasmodesmata, strigolactone biosynthesis, and xylem formation. Columns represent *Rafflesia*-infected and *Sapria*-infected conditions. Numerical values in each cell show the exact log2 fold change.

### Genes involved in gall formation

To investigate the presence of transcripts of genes that have previously been demonstrated in gall formation, we examined the expression patterns of phenylpropanoid and flavonoid biosynthesis genes identified in *Quercus infectoria* galls [14]. Using previously published datasets, we saw that several key biosynthetic genes showed strong upregulation relative to *Rafflesia speciosa* seeds [25] during floral bud development in *R. cantleyi* [26], (Figure 4). Most notably, leucoanthocyanidin dioxygenase 1 exhibited the highest upregulation in the floral bud stage (FBS2; log2FC = 7.94), followed by naringenin, 2-oxoglutarate 3-dioxygenase (log2FC = 6.10), chalcone synthase 2 (log2FC = 5.97), and flavonol synthase/flavanone 3-hydroxylase (log2FC = 4.93). In contrast, genes encoding early steps of the shikimate pathway were generally downregulated in infected tissues from the bud-root interface. This differs from the previous sampling approach where *R. cantleyi* bud-only samples at the FBS2 stage were analyzed [26], whereas this study examines both *Sapria* and *Rafflesia* buds at the FBS1 stage. Here, 3-deoxy-7-phosphoheptulonate synthase and 3-dehydroquinate synthase showed reduced expression in both *R. speciosa* and *S. himalayana* infected tissues (log2FC: -2.1 to -2.9). Notably, several genes involved in lignin biosynthesis were also downregulated in infected tissues, including hydroxycinnamoyltransferase 4, which showed the most dramatic downregulation in FBS2 (log2FC = -12.11), cinnamyl alcohol dehydrogenase 5 (log2FC: -2.5 to -2.8) and cinnamoyl-CoA reductase 1 (log2FC: -0.7 to -2.5). The expression patterns of cytochrome P450 genes varied considerably. While flavonoid 3’,5’-hydroxylase CYP75B138 and flavonoid 3’-monooxygenase showed strong downregulation in infected tissues (log2FC: -1.7 to -5.8), p-coumaroyl 3’-hydroxylase CYP98A-C2 exhibited upregulation in *S. himalayana* infected tissue (log2FC = 2.2) but downregulation in *R. speciosa* infected tissue (log2FC = -0.6). Interestingly, several genes showed dramatically different responses between the two parasite species, with flavonol synthase/flavanone 3-hydroxylase being strongly upregulated in *S. himalayana* infected tissue (log2FC = 6.6) but unchanged in *R. speciosa* infected tissue (log2FC = 0.0). These differential expression patterns suggest complex regulation of specialized metabolism during parasitic plant development and host infection.

**Figure 4:**
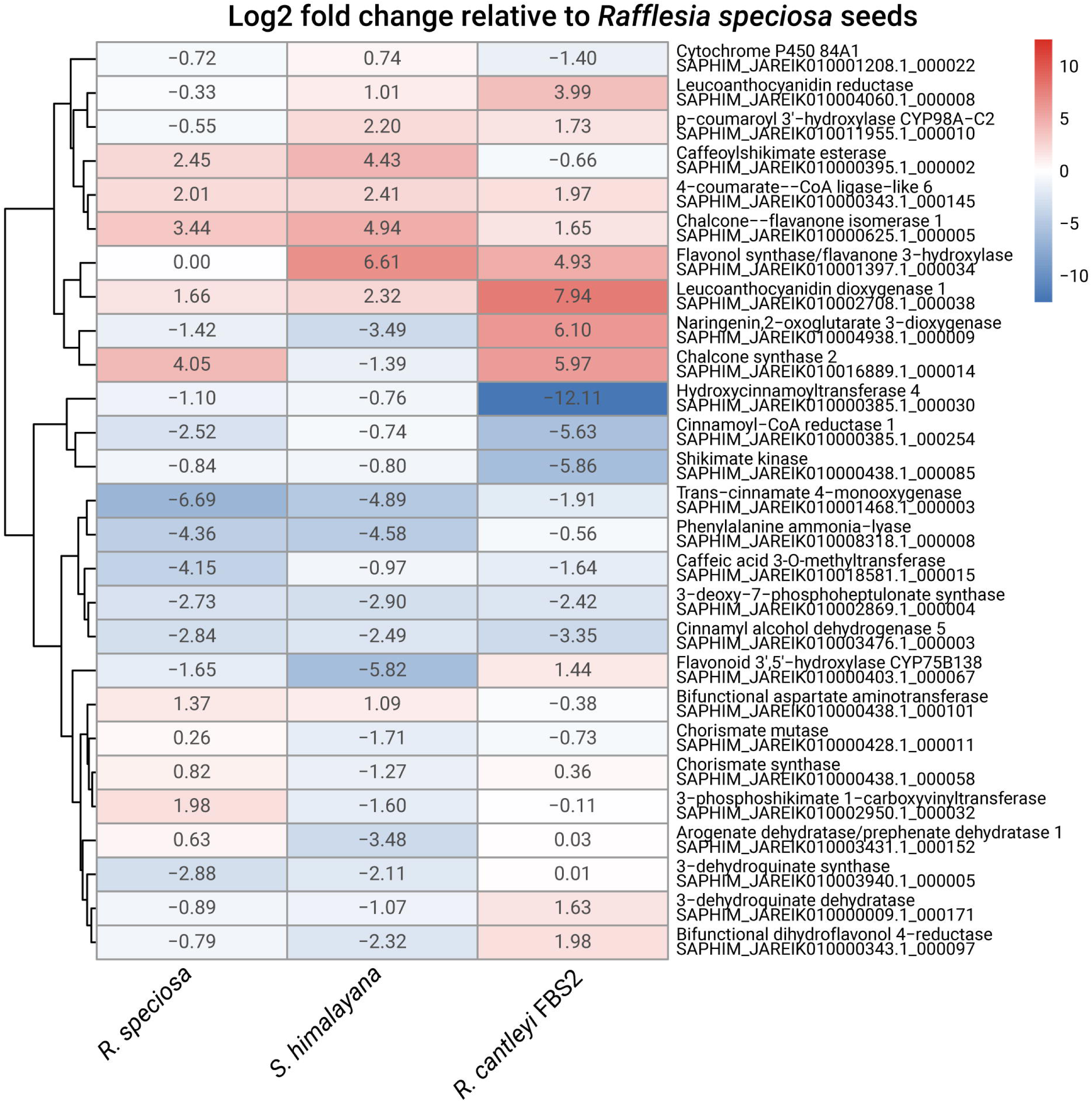
Expression patterns of phenylpropanoid and flavonoid biosynthesis genes previously identified in *Quercus infectoria* galls [14]. Heatmap shows log2 fold changes in gene expression relative to *Rafflesia speciosa* seeds across different parasitic plant samples: *R. speciosa*, *S. himalayana*, and *R. cantleyi* floral bud stage 2 (FBS2). Gene names and corresponding *Sapria* gene identifiers are shown on the right. Red indicates upregulation and blue indicates downregulation.

We then cross-referenced our datasets with a previously published set of thirty-eight genes upregulated in 4 different galls [16]. Several of these gall-associated genes showed strong differential expression in parasitic plants (Supplementary Figure 2). Most notably, genes encoding membrane-associated proteins displayed pronounced upregulation in both *Rafflesia* and *Sapria* infected tissues, including CYSTEINE-RICH TRANSMEMBRANE MODULE 4 (log2FC: 13.31 and 17.74, respectively) and cotton fiber protein (log2FC: 6.75 and 10.45, respectively). Peroxidase 25 also showed strong upregulation in both parasitic species (log2FC: 7.75 and 8.69). In contrast, several transcription factors exhibited tissue-specific expression patterns. While WRKY transcription factor 23 was upregulated in *Sapria* (log2FC: 3.57), it was downregulated in *Rafflesia* infected tissue and showed progressive downregulation through floral bud development (FBS1-3: -1.86 to -11.67). Similarly, heat stress transcription factor B-1 displayed initial upregulation in infected tissues but marked downregulation in later floral bud stages (FBS3: -10.93). Notably, genes involved in oxidative stress responses showed divergent patterns, with respiratory burst oxidase homolog protein C downregulated in *Rafflesia* infected tissue and FBS stages but upregulated in *Sapria*.

To investigate a possible bacterial origin in *Rafflesia*/*Sapria* gall formation, we compared our datasets against two sets of up- and downregulated genes from previously published transcriptome analysis of crown gall in radish [27]. The expression patterns showed largely inverse relationships between bacterial-induced crown galls and parasitic plant tissues (Supplementary Figure 3). Many genes upregulated in radish galls showed significant downregulation in parasitic plants. For example, XM_018620377 exhibited strong downregulation in both *R. speciosa* and *S. himalayana* infected tissues (log2FC: -8.06 and -4.84, respectively), while XM_018596275 showed consistent downregulation across infected tissues (log2FC: -2.57 and -3.04). Conversely, several genes that were downregulated in radish galls displayed strong upregulation in parasitic plants, particularly XM_018627109 (log2FC: 11.41 and 14.79 in *R. speciosa* and *S. himalayana*, respectively) and XM_018594057 (log2FC: 9.71 and 7.52). These contrasting expression patterns between bacterial-induced crown galls and parasitic plant tissues suggest that *Rafflesia*/*Sapria* gall formation likely occurs through distinct molecular mechanisms rather than bacterial manipulation, as the genetic signatures are largely opposite to those observed in crown galls.

In order to test if galls might be induced by the *Tetrastigma* host, we searched the *Tetrastigma magnum* genome for the presence of virulence factors known from previous studies in arthropods, such as effectors identified in the wheat pest *Mayetiola destructor* and gall wasps. However, when we BLAST-searched the *T. magnum* genome for these proteins, we only obtained low confidence hits with low sequence identity (below 40%). Only two exceptions were found: TETMAG_ptg000030l_000661.1, which matched the arthropod effector Mdes002782 with 67% identity but was most likely a heat shock protein, and TETMAG_ptg000003l_000044.1, which showed 60.7% identity with an oxidoreductase that is, however, also present in rice as locus Os09g0248900. The absence of clear homologs to known arthropod gall-inducing effectors in the *T. magnum* genome suggests that parasitic plant gall formation is not induced by the host in this case or occurs through different molecular mechanisms than those documented in insects.

We then analyzed fungal reads from parasitic plant-host interface tissues and found distinct taxonomic compositions between *Rafflesia*- and *Sapria*-infected *Tetrastigma* samples (Supplementary Figure 4). Of the entire dataset, we could assign 1.81% and 5.45% of reads to fungi in *Rafflesia*- and *Sapria*-infected samples, respectively (Supplementary Table 2). Within the fungal reads, we observed that *Ascomycota* dominated over *Basidiomycota* in *Rafflesia* (1.00% vs 0.21%), where we identified *Pyricularia grisea* (621k reads) and *Colletotrichum higginsianum* (105k reads) as prominent species. The *Sapria* sample exhibited higher proportions of both *Ascomycota* (2.93%) and *Basidiomycota* (1.65%), where we detected high abundances of *Sugiyamaella lignohabitans* (569k reads) and *Kwoniella botswanensis* (464k reads). We found substantial presence of *Rhizophagus irregularis* in both samples (2.01M reads in *Rafflesia*, 521k reads in *Sapria*). When these pre-filtered potential fungal reads were mapped to the *Rhizophagus irregularis* reference genome [28], the *Rafflesia*-infected sample yielded 25.21% (506,413 reads) uniquely mapped reads and 4.15% (83,364 reads) reads mapped to multiple loci, with a mismatch rate of 2.14%, while the *Sapria*-infected sample showed 26.73% uniquely mapped reads and 3.67% reads mapped to multiple loci, with a mismatch rate of 2.35%. Mapping to the *Pyricularia grisea* genome [29] (total reads: 620,931), yielded uniquely mapped reads of 12.56% (77,989 reads) and 18.77% for *Rafflesia*-infected and *Sapria*-infected samples, with multiple loci mapping rates of 3.33% (20,677 reads) and 6.87% and mismatch rates of 2.34% and 1.11%, respectively. Among the top expressed genes, a ubiquitin domain protein showed the highest expression in *Rafflesia*-infected *Tetrastigma*, while a Tubulin/FtsZ-like protein was highest in *Sapria*-infected *Tetrastigma*. Three genes were shared among the top 20 expressed genes: a histone protein, a ubiquitin domain protein, and the above-mentioned tubulin/FtsZ-like protein (Supplementary Figure 5). Although not among the top 20 expressed transcripts, we detected expression of several symbiosis-related genes: an MFS transporter had higher expression in *Sapria*-infected *Tetrastigma* (3521.2 TPM vs 61.7 TPM), and transcription factor Ste12 maintained high expression in both contexts (2949.6 TPM in *Sapria*, 1705 TPM in Rafflesia). An ammonium transporter showed higher expression in *Rafflesia*-infected *Tetrastigma* (880.34 vs 353.16 TPM). While this suggests arbuscular mycorrhizal associations at the host-parasite interface in established infections, it is notable that such associations were not observed in *Rafflesia* seeds [25].

Lastly, we investigated the top expressed genes from the pathogenic fungus *Pyricularia grisea*, whose reads were present in both our datasets (*Rafflesia* and *Sapria*-infected *Tetrastigma*). Our analysis revealed similar expression profiles with two genes dominating the expression landscape in both samples (Supplementary Figure 6): an ELMO domain-containing protein (PYRGRI_NC_044973.1_000642) and an uncharacterized gene (PYRGRI_NW_022156720.1_000292), both showing expression levels above 250,000 TPM. Several metabolic enzymes were also highly expressed, including O-methyltransferase COMT-type (>50,000 TPM) and anthranilate synthase (>17,000 TPM), which are involved in the modification of salicylic acid and biosynthesis of tryptophan, an auxin precursor, respectively. Other abundant transcripts included genes encoding proteins involved in oxidative stress responses (superoxide dismutase), cellular signaling (WD40-repeat containing proteins), and membrane transport (zinc/iron permease).

## Discussion

Our dual transcriptome analysis of *Tetrastigma* infected by *Rafflesia* and *Sapria* reveals distinct host responses to these closely related parasitic plants, providing insights into their potentially divergent infection strategies and host-parasite interactions.

### Comparative gene expression analysis in *Tetrastigma* hosts infected by *Rafflesia* vs. *Sapria*

The difference in the proportion of up- and down-regulated genes in *Tetrastigma*’s response to *Rafflesia* (88% up, 12% down) versus *Sapria* (98% up, 2% down) suggests different host-parasite dynamics. The strong upregulation trend in both cases indicates an active response by *Tetrastigma* to infection, rather than a generalized suppression of host gene expression. However, the near-complete dominance of upregulation in *Sapria*-infected *Tetrastigma* suggests a broader or more intense host response, which may reflect a more disruptive infection process or a less fine-tuned host–parasite interaction compared to the *Rafflesia* system. The nature of the most highly differentially expressed genes provides further insights. In *Rafflesia* infection, the upregulation of BURP domain proteins, zinc finger proteins, and glycoside sugar transporters suggests a reprogramming of host developmental processes. BURP proteins reinforce cell walls for abiotic stress tolerance [30], zinc finger proteins activate genes for antioxidants and osmoprotectants [31], and glycoside sugar transporters allocate sugars for energy and defense during pathogen attacks and key developmental stages [32]. The concurrent upregulation of these genes could indicate an active defense strategy, where sugars act as metabolic fuel, zinc fingers activate defense genes, and BURP proteins strengthen cell walls. The downregulation of defense-related genes like LRR and TIR domain proteins indicates a suppression of host immune responses by *Rafflesia*. In contrast, *Sapria* infection upregulates EARLY NODULIN 93 (ENOD93) and glutathione S-transferase (GST) genes, with ENOD93 promoting symbiotic nitrogen fixation [33] and GST mitigating oxidative damage. Meanwhile, the downregulation of PPR proteins impairs mitochondrial and chloroplast functions [34], and reduced cytochrome P450 enzyme expression limits secondary metabolite and phytohormone biosynthesis, prioritizing survival over growth and defense [35].

The COG analysis supports the observation of distinct host responses to Rafflesia and Sapria infection. Both parasites induced upregulation across most functional categories; however, the magnitude was generally greater in Rafflesia infection, particularly in signal transduction and defense mechanisms – hallmarks of an active host response to biotic stress [36,37]. This may reflect a more focused or efficient reprogramming of host processes by Rafflesia. In contrast, Sapria infection produced a more balanced pattern of up- and downregulation across several categories, including secondary metabolism and defense, suggesting a more complex or variable interplay between host immunity and parasite manipulation. Notably, genes associated with posttranslational modification were highly upregulated in Sapria-infected roots, potentially facilitating a more dynamic and rapid immune response [38]. GO enrichment further revealed that Sapria infection broadly reprograms host cellular functions, with significant enrichment in vesicle trafficking, protein turnover, and stress-response pathways. This pattern may indicate either widespread host stress or an expansive remodeling of cellular infrastructure to accommodate the parasite. By contrast, Rafflesia infection resulted in only a single significant GO term (cytosolic ribosome) suggesting a more targeted or constrained manipulation of host functions that may avoid broad systemic disturbance.

The analysis of specific gene expression patterns in *Tetrastigma* reveals nuanced differences in host responses to *Rafflesia* and *Sapria*, and the results highlighted two main perspectives: parasitic-induced physical stress response by the host (as shown on the cell wall integrity genes and xylem differentiation) and host immune response (as shown on the plant immune response, strigolactone production, and plasmodesmata-related genes). The differential regulation of cell wall integrity genes, such as the strong upregulation of *THE1, FER* and *MCA1* during *Sapria* infection, suggests distinct impacts on host cell structure. THE1 acts as a surveillance protein for cell wall defects, FER regulates cell-cell communication [39] and MCA1 functions as a mechanosensory protein involved in jasmonic acid production during cell wall remodeling [40]. The elevated expression of these genes in *Sapria*-infected roots may indicate higher stress levels due to the comparatively larger swelling of the parasite bud relative to host root size, as opposed to the *Rafflesia* bud.

The upregulation of plant-plant interaction genes like *Or7* in both infections, but with different magnitudes, indicates a common but parasite-specific recognition response. This protein is an LRR receptor-like kinase found in *Orobanche cumana* that serves as a parasitic resistance protein [41], thus the high expression, and interestingly, implying a possible homologous protein expression that also occurs in *Tetrastigma*. Similarly, SOBIR1 protein, a protein involved in plant immune response [42] showed a high amount of expression in the *Sapria*-infected *Tetrastigma* root.

Interestingly, both infections result in the downregulation of most strigolactone biosynthesis genes, including CCD8, MAX1, and PDR1, which may represent a parasite strategy to manipulate host development and branching [43]. Contrasting regulation of xylem formation genes further emphasizes differences in parasite strategies: KNAT7 and SND1, involved in secondary wall biosynthesis [44–47], are upregulated in *Sapria* but not in *Rafflesia*. In contrast, VND6 and VND7, key regulators of xylem differentiation [47,48], are downregulated in *Sapria* but upregulated in *Rafflesia*. These differences likely stem from variations in the age and size of the host roots infected by the two parasites.

Plasmodesmata-related genes also exhibit notable differences. The upregulation of CalS1, which enhances callose deposition and cell wall defense [49], and BAM, which assists siRNA movement [50] in *Sapria*-infected roots, points to a higher stress response in these hosts. While the possibility of interspecific plasmodesmata between host and parasite remains unclear, the expression fluctuations of these genes suggest potential cellular connections facilitating parasite-host interactions, resembling structural support mechanisms in arthropod-induced galls [51]. Similarities include increased defense-related compounds, such as tannins, flavonols, and salicylates, in outer tissues [52], suggesting parallel strategies for maintaining structural integrity during parasitism.

### Distinct gene expression profiles in the parasites *Rafflesia speciosa* and *Sapria himalayana*

The high expression of cytochrome b/b6 and cytochrome c oxidase in both *Rafflesia* and *Sapria* reflects their elevated energy demands and metabolic adaptation to parasitic lifestyles. As integral components of the mitochondrial electron transport chain, these proteins facilitate efficient ATP production, supporting the energy-intensive processes of floral development and host resource extraction. The shared expression of NADH:ubiquinone oxidoreductase subunits and ATP synthase complex components further underscores the critical importance of oxidative phosphorylation in both parasites. However, the dramatically higher expression levels of these mitochondrial genes in *Rafflesia* compared to *Sapria* suggests distinct parasitic strategies. *Rafflesia*’s exceptionally high expression of energy production machinery indicates an emphasis on rapid and intensive resource acquisition from the host. In contrast, *Sapria*’s relatively lower but still substantial expression of these same genes may reflect a more measured approach to energy metabolism. The presence of leucine-rich repeat domain proteins in both species, albeit at vastly different expression levels (*Rafflesia*: 17,737 TPM vs. *Sapria*: 7 TPM), suggests both parasites maintain some capacity for host recognition or defense responses, though *Rafflesia* appears to invest much more heavily in these processes. This differential expression pattern indicates that while both parasites rely on similar fundamental metabolic machinery, they have evolved distinct strategies for host interaction and resource utilization. Though both *Rafflesia* and *Sapria* show KEGG enrichment in mRNA surveillance, RNA transport, and RNA degradation pathways highlighting the parasites’ ability to dynamically regulate its transcriptome and maintain RNA integrity, they also showed differences. *Rafflesia*’s enrichment in amino acid biosynthesis and energy production pathways aligns with its gene expression profile, suggesting a parasitic strategy focused on efficient nutrient extraction and utilization. *Sapria*’s enrichment in pathways related to membrane biogenesis, oxidative stress response, and immune signaling indicates a more complex interaction with the host, possibly involving more extensive tissue invasion or manipulation of host defenses.

Our study reveals that *Rafflesia* and *Sapria*, despite their close evolutionary relationship, induce markedly different transcriptional responses in their *Tetrastigma* hosts, *T. magnum* and *T. obovatum*, respectively. These differences span broad functional categories and specific pathways, suggesting distinct infection strategies and host-parasite dynamics. *Rafflesia* appears to induce a strong but potentially more targeted reprogramming of host processes, while *Sapria* triggers a broader, more intense host response. This suggests that *Rafflesia* has a more precise mechanism for manipulating host resources or responses, possibly minimizing collateral effects on the host’s overall physiology. In contrast, *Sapria* seems to elicit a more widespread and intense transcriptional response from the host. This could mean that *Sapria* affects a larger array of host pathways or triggers a more systemic reaction, such as heightened stress responses or broader immune activation, likely imposing greater overall stress on the host. The distinct parasitic strategies of *Rafflesia* and *Sapria* have important implications for host physiology, which could inform horticultural practices and *ex situ* conservation efforts. Understanding the molecular basis of these interactions could enable targeted manipulation of host signaling pathways to enhance compatibility with either parasite. *Rafflesia*’s ability to modulate specific pathways might make it easier to establish long-term parasitic associations with minimal disruption to host physiology. *Sapria*’s induction of a more intense immune response and stress pathways suggests a need for careful management of host conditions, as prolonged stress could reduce host longevity and reproductive output. In addition, these findings open up several avenues for future research. Comparative metabolomic analyses could provide insights into how the observed gene expression changes affect nutrient flow between host and parasite. Finally, evolutionary analyses comparing these patterns across multiple Rafflesiaceae species could shed light on the diversification of parasitic strategies within this fascinating plant family.

### Molecular parallels between Rafflesiaceae parasitism and gall formation: A potential role for fungi?

Our analyses also show that Rafflesiaceae parasitism shares molecular signatures with gall formation, while suggesting a mechanism distinct from both bacterial and insect-induced galls. The expression patterns of phenylpropanoid and flavonoid biosynthesis genes in Rafflesiaceae closely mirror those observed in *Quercus infectoria* galls [14], particularly during floral bud development. Across Rafflesiaceae samples, the strong upregulation of leucoanthocyanidin dioxygenase and flavonol synthase, which contribute to the synthesis of flavonoids and tannins and are essential for gall formation, highlights their roles in chemical defense and structural reinforcement, possibly facilitating the development of the extensive interface tissue characteristic of these parasites. Interestingly, the concurrent downregulation of hydroxycinnamoyltransferase 4 and cinnamoyl-CoA reductase in Rafflesiaceae samples suggests a shift away from lignin biosynthesis, favoring chemical defenses and tissue modifications. Consistent with this, prior metabolomic studies on *Rafflesia* buds [15] revealed an abundance of gallic acid derivatives, further supporting the idea that *Rafflesia* buds share functional similarities with galls, utilizing comparable metabolic pathways to adapt to their parasitic lifestyle.

Our data suggest a potential role for fungi, particularly the hemibiotrophic pathogen *Pyricularia grisea*, in facilitating gall formation. The high expression of *P. grisea* genes involved in hormone manipulation, such as O-methyltransferase, which may alter salicylic acid levels [36], and anthranilate synthase (a precursor for auxin biosynthesis), indicates that these fungi might inadvertently create conditions conducive to gall induction. While *P. grisea* is primarily known as a foliar pathogen in grasses, it also inhabits the rhizosphere in natural environments [53]. The consistent presence and similar expression patterns of *P. grisea* in both *Rafflesia* and *Sapria* systems suggest this association may not be coincidental, potentially representing a novel dimension in the evolution of plant parasitism. Additionally, the detection of substantial populations of mycorrhizal fungus *Rhizophagus irregularis* at the host-parasite junctions in *Rafflesia* and *Sapria* suggests an involvement of fungal symbiotic activity in *Rafflesia*- and *Sapria*-infected *Tetrastigma*, with certain core cellular programs active in both samples with three overlapping genes in the top 20 – histone, ubiquitin, and tubulin. Interestingly, mycorrhizal taxa were not observed in *Rafflesia* seeds [25]. The presence of MFS transcripts is consistent with known roles of MFS transporters in nutrient exchange during fungal-plant symbiosis [54]. Ste12’s expression in both samples indicates conserved regulation of phosphate homeostasis, similar to its role in other mycorrhizal interactions [55]. Finally, the ammonium transporter’s higher expression in *Rafflesia*-infected *Tetrastigma* suggests different nitrogen transport dynamics between hosts, reflecting the known importance of these transporters in fungal-plant nitrogen exchange [56]. This dynamic echoes the tripartite interaction observed in the holoparasite *Cytinus hypocistis* and its Cistaceae host, both colonized by mycorrhizae. It has been suggested that arbuscular mycorrhizae could act as gall inducers, triggering root cell expansion through gibberellin signaling [57]. Microscopic analyses of the Rafflesiaceae-host junction are warranted to confirm these associations and explore the role of microbial interactions in gall-like structures.

Together, these findings reveal that the Rafflesiaceae symbiotic system is a complex ecosystem of plant-microbial interactions, extending far beyond the visible host-parasite interface.

### Limitations and future directions

While this study offers novel insights into Rafflesiaceae parasitism and host responses, several limitations remain. The lack of biological replicates, due to the rarity of infected material, limits the statistical resolution of our differential expression analysis, despite compensation through deep sequencing and stringent filtering. Future work should prioritize replication across host-parasite pairs and developmental stages. Cross-species comparisons based on TPM values introduce biases due to differences in gene models and transcript lengths, and ortholog-based analyses would improve cross-species resolution. Additionally, the gall-like traits and microbial associations we identify remain correlative. Functional validation through microscopy, in situ assays, or gene knockouts in model systems will be crucial to test these hypotheses. Finally, we provide a reference genome for *Tetrastigma magnum* and the first dual transcriptomic dataset for Rafflesiaceae. Integration with metabolomic and proteomic data will be necessary to fully characterize the biochemical and ecological complexity of these unique parasitic systems. These integrative efforts will be key to advancing our understanding of parasitic plant biology and informing effective conservation strategies for Rafflesiaceae.

## Materials and Methods

### Plant samples

Samples of *Sapria himalayana* and its host *Tetrastigma cauliflorum* were obtained from Queen Sirikit Botanic Garden area, Mae Rim, Chiang Mai, Thailand in January 2023. Samples of *Rafflesia speciosa* and its host *Tetrastigma magnum* were obtained from the region of Miagao, Iloilo Province, Panay Island, Philippines in August 2023. The infected samples (Supplementary Figure 7) were collected from the flower bud–host root interface, specifically the proximal region of the Rafflesiaceae bud cupule where it directly intersects with the *Tetrastigma* root tissue. Uninfected *Tetrastigma* tissues (*T. cauliflorum* and *T. magnum*) were also obtained using a biopsy tubular blade for comparison to infected tissues. All samples were immediately placed in microtubes pre-filled with DNA/RNA Shield™ (Zymo Research) to preserve nucleic acids during transit. The uninfected root of *T. cauliflorum* used in this study was also processed, sequenced, and used in a previous miRNA identification study [58]. Due to the rarity of these samples, no replicate was obtained, but the depth of the sequencing was increased to ensure the inclusion of as many reads as possible. For genome sequencing of *Tetrastigma magnum*, fresh leaf material from the United States Botanic Garden (USBG) cultivar 2017-0203 (originally sourced from Miagao, Iloilo, Philippines) was used. All necessary collecting permits were obtained from the appropriate authorities (National Research Council of Thailand and Philippine Department of Natural resources).

### RNA extraction and sequencing

The samples were homogenized and had the total RNA extracted using standard protocols. The results of total RNA samples were then converted into cDNA. Total cDNA samples were sequenced by GENEWIZ (New Jersey) using paired-end sequencing on an Illumina NovaSeq 6000 platform.

### Genomic DNA extraction

A single leaf (1 g) of *Tetrastigma magnum* USBG cultivar 2017-0203 was processed to isolate nuclei using the Circulomics Nuclei Isolation – LN2 Plant Tissue Protocol (Document ID: NUC-LNP-001). Subsequently, 30 μg of high molecular weight genomic DNA was extracted from these nuclei using the Nanobind HMW DNA Extraction – Plant Nuclei Protocol (Document ID: EXT-PLH-001), as part of the Nanobind Plant Nuclei Big DNA Kit (PacBio).

### PacBio HiFi genomic sequencing

Genomic DNA was quantified on a Qubit 3.0 and sized using a Genomic ScreenTape on an Agilent TapeStation 4200. Samples that met QC guidelines (50% >30 kb) were taken into library preparation. DNA was sheared using a Covaris g-Tube and centrifuged at 2,750 rpm in an Eppendorf 5430R centrifuge to target 13-20 kb fragments. The DNA underwent 4 full passes prior to cleanup with SMRTbell cleanup beads. This DNA was then used for generating SMRTbell libraries using the PacBio SMRTbell 3.0 kit according to manufacturer protocol. Libraries were size-selected using 35% Ampure PB beads to remove fragments <5 kb. Final libraries were quantified using a Qubit 3.0 and sized using an Agilent Femto Pulse with a Genomic DNA 165 kb kit. Final libraries were loaded onto the PacBio Revio according to manufacturer protocol. The targeted loading concentration was between 200-300 pM per sample. Sequencing happened for 30 hours per sample.

### Genome assembly and annotation

The *T. magnum* genome was assembled using Hifiasm 0.16.0 [59] with PacBio HiFi reads, with options for primary contig generation. Gene prediction of all assemblies used in this study was carried out using Helixer v0.3.3 [60], with the land plant lineage as reference. The resulting GFF3 file was processed using gffread [61] to extract protein sequences from the predicted genes and used for functional annotation with InterProScan 5.55-88.0 [62] and EggNOG-mapper 2.1.12 [63]. The completeness of the genome assembly and gene set was assessed using Benchmarking Universal Single-Copy Orthologs (BUSCO) 5.3.2 [64] analysis against the eudicots_odb10 lineage dataset. The assembly and annotation of the *Tetrastigma magnum* genome have been deposited in the NCBI GenBank under BioProject PRJNA1255605 and genome accession number JBNLLH000000000.

### RNA-seq data mapping

The genome of *Tetrastigma magnum* and *Sapria himalayana* were indexed using STAR v2.7.3a [65] with default parameters. The resulting BAM files were processed using featureCounts v2.0.1 [66] with options to generate read counts per gene.

This analysis pipeline was applied to investigate host gene expression responses to parasitic plant infection. For the *Rafflesia* infection analysis, uninfected *Tetrastigma magnum* (a host of *Rafflesia*) was compared to *Rafflesia*-infected *T. magnum*. For the *Sapria* infection analysis, uninfected *Tetrastigma cauliflorum* (a host of *Sapria*) was compared to infected *Tetrastigma obovatum*, also a host of *Sapria*. For these comparisons, the transcriptomic data of uninfected samples were mapped to the reference genome of *Tetrastigma magnum*. For the infected samples, data were separately mapped to the reference genomes of *Tetrastigma magnum* and *Sapria himalayana* to distinguish host and parasite gene expression. The raw RNA-seq reads generated in this study have been deposited in the NCBI Sequence Read Archive (SRA) under BioProject accession number PRJNA1215969.

### Differential Expression Analysis

Differentially expressed gene (DEG) analysis was conducted with edgeR [67] running in RStudio 2024.09.0 Build 375 using R version 4.4.1 (Posit PBC). Data was loaded using readr processed with edgeR, and visualized using dplyr [68]. Gene expression levels were calculated as Transcripts Per Million (TPM) using custom functions. Annotation data was extracted and processed using the stringr package. Differential expression analysis results were visualized as heatmaps using pheatmap [69], with plot arrangement facilitated by gridExtra and grid. Comparison of differentially expressed genes between *Rafflesia* and *Sapria* infections was performed using dplyr and visualized with proportional Venn diagrams using eulerr. Additional visualizations were created using ggplot2, with multi-panel plots arranged using patchwork. Data transformations and formatting were aided by tidyr [70] and scales. This analysis pipeline was applied to *Tetrastigma magnum* samples infected with *Rafflesia* and *Sapria* to compare host gene expression responses to these parasitic plants.

### COG, GO, and KEGG functional analyses

Additional functional analyses were performed also using RStudio 2024.09.0 Build 375 running R version 4.4.1 (Posit PBC). The analyses on *Tetrastigma* infected with *Rafflesia* and *Sapria*, included annotations of Clusters of Orthologous Groups (COG) to compare differentially expressed genes, Gene Ontology (GO) enrichment analysis was performed using R to compare differentially expressed genes that belong to specific GO groups (e.g. biological process or BP, cellular component or CC, and metabolic function or MF), and Kyoto Encyclopedia of Genes and Genomes (KEGG) pathway enrichment analysis.

Both COG and GO data were obtained from the *Tetrastigma magnum* annotation data, while for KEGG data annotations, the input was acquired from eggNOG-mapper output. Data editing and filtering were conducted using dplyr package for COG, and for GO and KEGG, data editing, filtering, and preprocessing were conducted using dplyer, tidyr, and stringr packages. GO terms were extracted and categorized into CC, MF, and BP types. The clusterProfiler package was used to perform GO enrichment analysis for both *Rafflesia* and *Sapria* infected samples, with a p-value cutoff of 0.05 and q-value cutoff of 0.2. The clusterProfiler package was used to perform KEGG enrichment analysis for both *Rafflesia* and *Sapria* infected samples, with a *p*-value cutoff of 0.05 and Benjamini-Hochberg adjustment for multiple testing. Visualization of COG distributions (with each DEG plotted on its log2fold change and assigned COG category, and the resulting plot were arranged with gridExtra and cowplot packages), GO terms comparison bar plot, and KEGG pathway enrichment rich factors were visualized using ggplot2 package.

### Pathway-Specific Gene Expression Analysis

The pathway-specific genes were picked on specific processes that were hypothetically involved with the impact of infection to the host plants. First pathway is the strigolactone production pathway to see if *Tetrastigma* might produce strigolactone that induce seed germination in other parasitic plants, like broomrapes (Orobanchaceae) [43]. Then, cell wall integrity proteins, which are involved in plant stress [71], and as Rafflesiaceae infection could hypothetically induce a certain stress to the host, the expression patterns of these genes could help to elaborate the host stress at the cellular level. Expression of proteins involved in cell-to-cell plasmodesmata formation were also observed, as it was predicted that parasite-host connection might induce interspecific cell plasmodesmata formation [49,72]. As Rafflesiaceae enters generative stage and the bud forms from the parasitic parenchymal endophyte, a previous study refers that host xylem formation was distorted by the parasitic growth [1], this group of xylem differentiation genes [73,74] were selected. Additionally, host immune response proteins in *Cuscuta* infection [19] were also checked, in case of possible homologous genes existence in *Tetrastigma*. The list of genes is available in Supplementary Table 3.

Pathway-specific gene expression analysis was performed using RStudio 2024.09.0 Build 375 running R version 4.4.1 (Posit PBC) to compare expression levels of selected genes in *Tetrastigma* under uninfected conditions and when infected with *Rafflesia* and *Sapria*. Gene information was extracted from FASTA files using stringr. Count data for each condition was processed using dplyr and tidyr. The edgeR 4.0 package [67] was used to create a DGEList object and calculate normalization factors, thereby normalizing gene expression levels across samples. Counts per million (CPM) values were then computed based on the normalized data.

## Acknowledgements

We would like to thank the Stem Cell Genomics Core at the Sanford Stem Cell Institute for providing sequencing services. J.M. sincerely thanks her collaborators in the Philippines, including the field guides and staff of Miag-ao municipality, with special acknowledgment to Mayor Richard Garin and Dr. Macario Napulan. Deep appreciation is also extended to Ronniel Pedales, Julie Barcelona, Marites Muyong and her family, and the dedicated team from the Philippine DENR. We are equally grateful for the support from Thailand’s NRCT and QSBG, the Philippine Bureau of Plant Industry, the U.S. Botanic Garden, USDA, and the research office at Pace University. Special thanks goes to Ari Novy, Jim Westwood, Soyon Park, Joseph Morin, Adrian Tobias, Thomas Lipscomb, Ghea Putri Cristy, Stephen Elliott, Hans Bänziger, Piyakaset Suksathan, and the late Wattana Tanming for their invaluable support. We acknowledge the contributions of the late Joanne Chory to this work, including funding. J.C. was an investigator of the Howard Hughes Medical Institute.

## Author Contributions

J.M. conceived the study. All authors analyzed the data and wrote the manuscript.

## Data Availability Statement

The assembly and annotation of the *Tetrastigma magnum* genome have been deposited in the NCBI GenBank under BioProject PRJNA1255605 and genome accession number JBNLLH000000000. The raw RNA-seq reads generated in this study have been deposited in the NCBI Sequence Read Archive (SRA) under BioProject number PRJNA1215969.

