## Supplementary Table 1 for "Deciphering Parasitic Strategies: Dual Transcriptomics Reveal Distinct Infection Mechanisms and Gall-like Traits in Rafflesiaceae"

**Supplementary Table 1: Genome assembly statistics and BUSCO assessment of *Tetrastigma magnum*.** The table shows genome size, GC content, predicted genes (CDS), and BUSCO completeness metrics. BUSCO analysis was performed using the eudicot dataset (eudicots_odb10) as reference. CDS: coding sequences.

| Assembly size [bp] | 2,376,545,202 |
| --- | --- |
| Number of contigs | 317 |
| Contig N50 [bp] | 75,267,427 (n=13) |
| Contig N90 [bp] | 24,534,822 (n=33) |
| Percent GC | 38.91 |
| Number of CDS | 47,190 |
| Complete BUSCOs | 94.4% |
| Single-copy BUSCOs | 89.8% |
| Duplicated BUSCOs | 4.6% |
| Fragmented BUSCOs | 3.5% |
| Missing BUSCOs | 2.1% |
