## Supplementary Table 2 for "Deciphering Parasitic Strategies: Dual Transcriptomics Reveal Distinct Infection Mechanisms and Gall-like Traits in Rafflesiaceae"

**Supplementary Table 2: RNA-seq mapping statistics for host and fungal reference genomes.** For fungal mappings, reads were pre-filtered by applying Kraken2 taxonomic classification to the original BAM files, from which new BAM files containing only the fungal-classified reads were created for subsequent mapping.

| **Reference Genome** | **Sample** | **Total Reads** | **Uniquely Mapped (%)** | **Uniquely Mapped (reads)** | **Multi-mapped (%)** | **Multi-mapped (reads)** | **Mismatch Rate (%)** |
| --- | --- | --- | --- | --- | --- | --- | --- |
| *T.*  *magnum*  (this study) | Uninfected  *T. magnum* | 191,919,296 | 68.85 | 132,136,935 | 9.36 | 17,963,646 | 0.42 |
|  | *Rafflesia*-infected  *T. magnum* | 376,724,846 | 42.81 | 161,276,507 | 6.72 | 25,316,310 | 0.36 |
|  | Uninfected  *T. cauliflorum* | 181,320,714 | 33.96 | 61,576,515 | 3.55 | 6,436,885 | 3.03 |
|  | *Sapria*-infected *T. obovatum* | 334,651,464 | 26.76 | 89,552,732 | 3.12 | 10,441,126 | 2.23 |
| *S. himalayana*  (Guo et al., 2023) | *Sapria*-infected *T. obovatum* | 334,651,464 | 38.55 | 129,008,139 | 3.08 | 10,307,265 | 0.41 |
|  | *Rafflesia*-infected  *T. magnum* | 376,724,846 | 31.45 | 118,479,964 | 10.13 | 38,162,927 | 3.12 |
|  | *Rafflesia speciosa* seeds  (Molina et al., 2023) | 172,545,511 | 19.52 | 33,680,884 | 0.54 | 931,745 | 4.50 |
|  | *Rafflesia cantleyi* FBS1  (Amini et al., 2017) | 11,626,735 | 20.03 | 2,328,576 | 3.91 | 454,712 | 4.86 |
|  | *Rafflesia cantleyi* FBS2  (Amini et al., 2017) | 8,956,135 | 19.10 | 1,710,957 | 1.65 | 147,635 | 5.07 |
|  | *Rafflesia cantleyi* FBS3  (Amini et al., 2017) | 11,269,613 | 20.41 | 2,300,490 | 1.71 | 192,291 | 5.09 |
| *R.*  *irregularis*  (Manley et al., 2023) | *Rafflesia*-infected  *T. magnum* (pre-filtered) | 2,008,778 | 25.21 | 506,413 | 4.15 | 83,364 | 2.14 |
|  | *Sapria*-infected *T. obovatum* (pre-filtered) | 520,711 | 26.73 | 139,186 | 3.67 | 19,110 | 2.35 |
| *P.*  *grisea*  (Gomez Luciano et al., 2019) | *Rafflesia*-infected  *T. magnum* (pre-filtered) | 620,931 | 12.56 | 77,989 | 3.33 | 20,677 | 2.34 |
|  | *Sapria*-infected *T. obovatum* (pre-filtered) | 664,110 | 18.77 | 124,653 | 6.87 | 45,624 | 1.11 |
