## Supplementary Table 3 for "Deciphering Parasitic Strategies: Dual Transcriptomics Reveal Distinct Infection Mechanisms and Gall-like Traits in Rafflesiaceae"

**Supplementary Table 3:** Selected pathway-specific genes.

| Pathway | Genes | References |
| --- | --- | --- |
| Strigolactone biosynthesis | *DWARF27, CCD7, CCD8, LBO, MAX1, PDR1* | (Cochetel et al., 2018) (Cochetel et al., 2018) |
| Cell wall integrity | *THE1, FEI2, FER, MCA1, WAK* | (Baez et al., 2022) (Baez et al., 2022) |
| Plasmodesmata formation | *BAM1, BAM2, CALS1, CALS8, PDLP5, PDLP7* | (Fischer et al., 2021) (Fischer et al., 2021)  (Iswanto et al., 2022) (Iswanto et al., 2021) |
| Xylem formation | *KNAT7, SND1, VND6, VND7* | (Hirai et al., 2021) (Hirai et al., 2021)  (Agusti and Blazquez, 2020) (Agusti and Blázquez, 2020) |
| Host immune response in *Cuscuta* | *Or7, SERK, SOBIR1, VuNB-LRR* | (Albert et al., 2021) (Albert et al., 2021) |
