## Supplementary Figure 1 for "Deciphering Parasitic Strategies: Dual Transcriptomics Reveal Distinct Infection Mechanisms and Gall-like Traits in Rafflesiaceae"

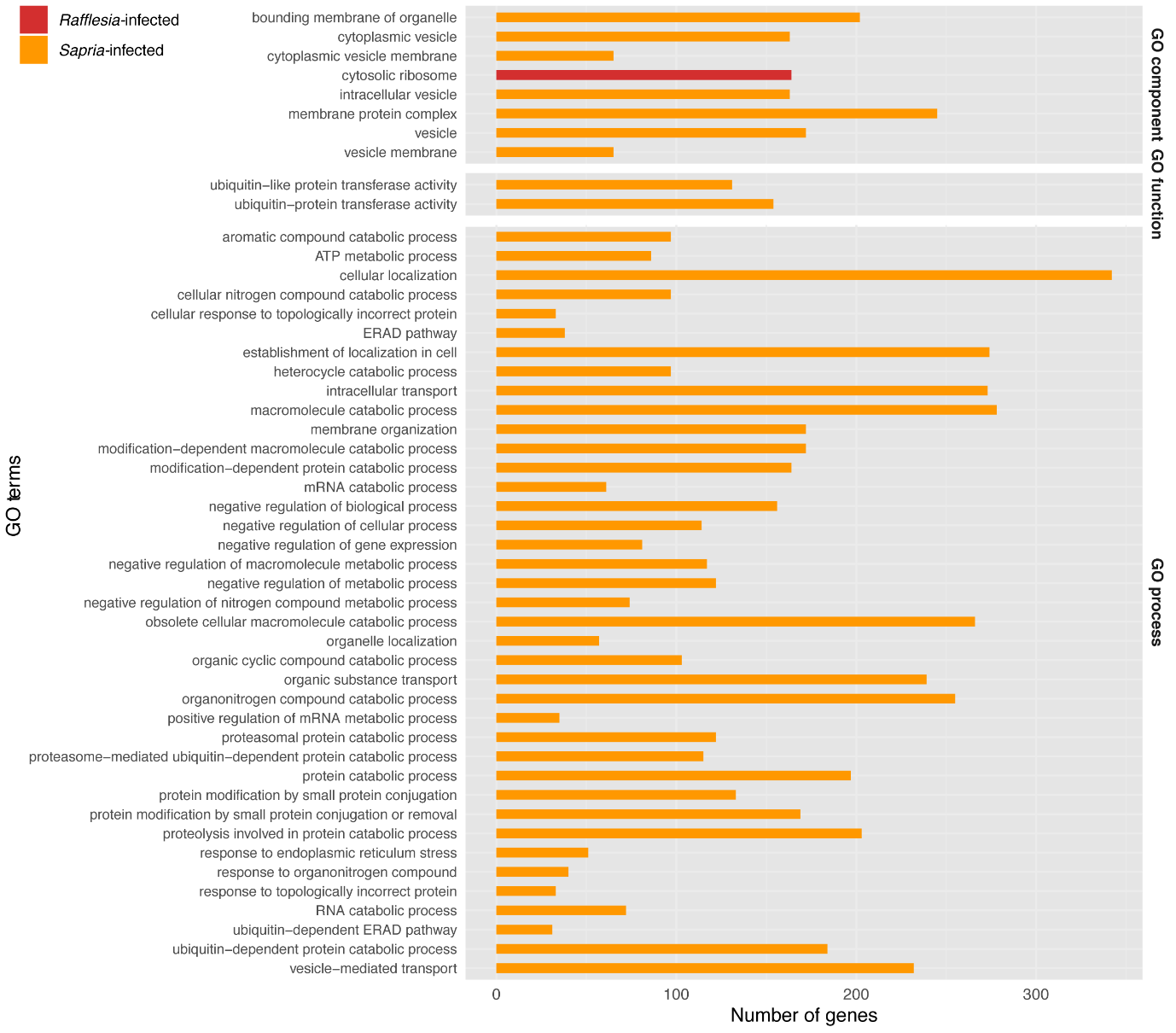

**Supplementary Figure 1: Comparison of GO (Gene Ontology) term enrichment between *Tetrastigma* infected by *Rafflesia* vs *Sapria*.** The bar plot shows the top enriched GO terms across three categories: cellular component, molecular function, and biological process. Each bar represents the number of genes associated with a particular GO term, with red bars indicating *Rafflesia* infection and orange bars indicating *Sapria* infection. The y-axis lists the GO terms, while the x-axis shows the number of genes.
