## Supplementary Figure 2 for "Deciphering Parasitic Strategies: Dual Transcriptomics Reveal Distinct Infection Mechanisms and Gall-like Traits in Rafflesiaceae"

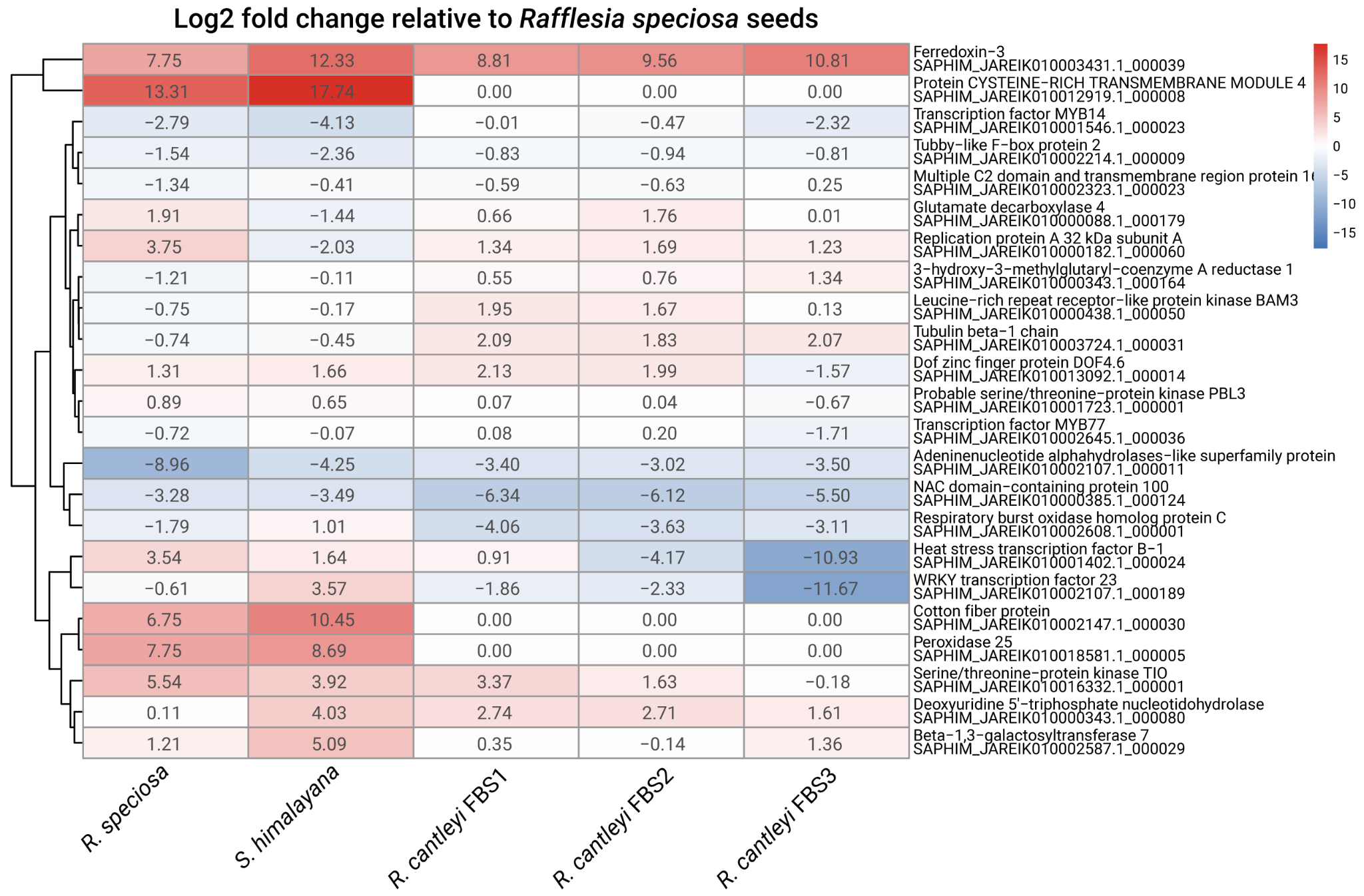

**Supplementary Figure 2: Expression patterns of gall-associated genes identified from comparative transcriptome analysis of diverse plant galls in (Takeda et al., 2019).** Heatmap shows log2 fold changes in gene expression relative to *Rafflesia speciosa* seeds across different parasitic plant samples: *R. speciosa*, *S. himalayana*, and three *R. cantleyi* floral bud stages (FBS1-3). Gene names and corresponding *Sapria* gene identifiers are shown on the right. Red indicates upregulation and blue indicates downregulation.
