## Supplementary Figure 3 for "Deciphering Parasitic Strategies: Dual Transcriptomics Reveal Distinct Infection Mechanisms and Gall-like Traits in Rafflesiaceae"

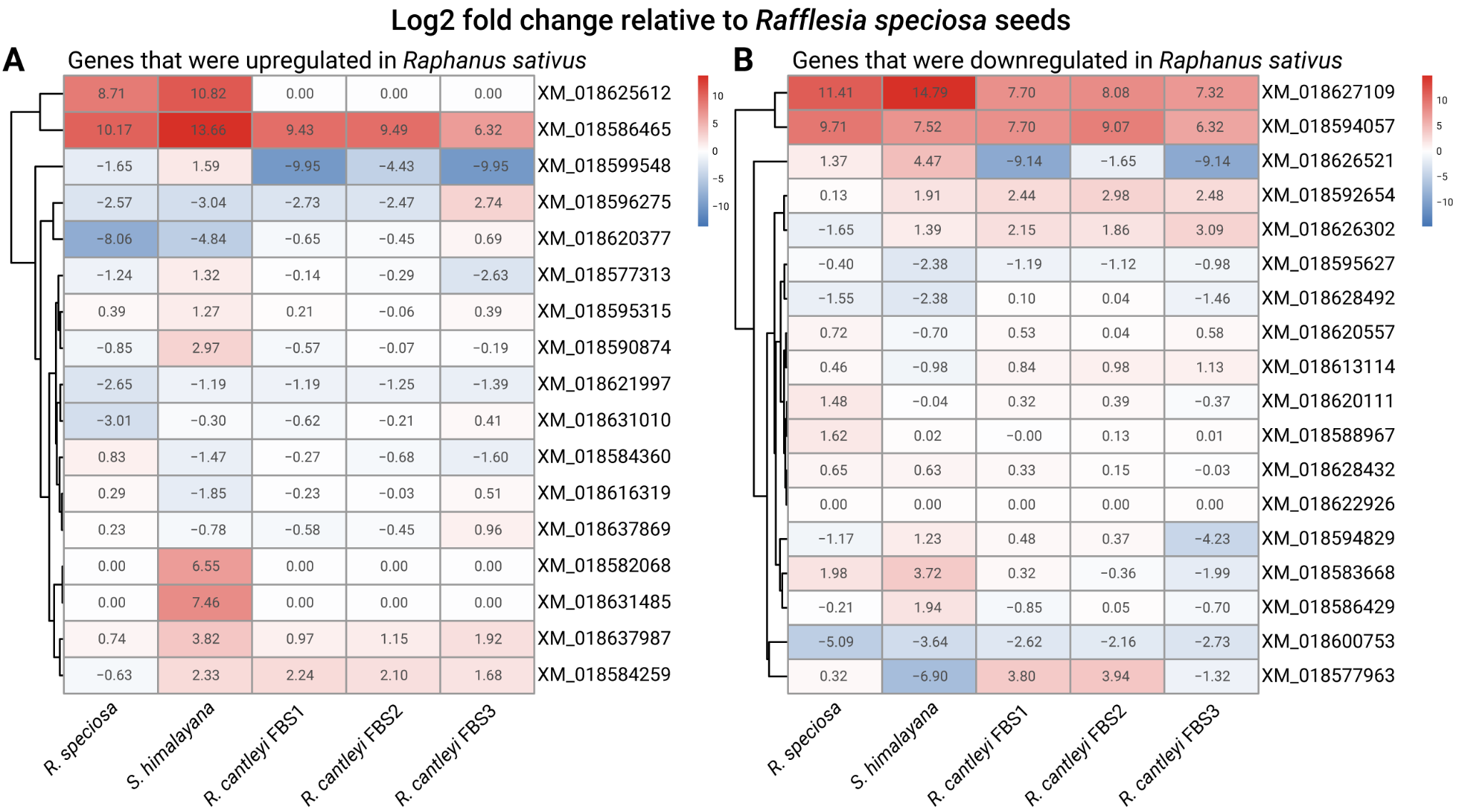

**Supplementary Figure 3: Expression patterns of genes previously identified as differentially regulated in crown gall in Raphanus sativus in (Tkachenko et al., 2021).** Heatmaps show log2 fold changes in gene expression relative to *Rafflesia speciosa* seeds for genes that were **A:** upregulated or **B:** downregulated (B) in *R. sativus* galls. Expression is compared across *R. speciosa*, *S. himalayana*, and three *R. cantleyi* floral bud stages (FBS1-3). Red indicates upregulation and blue indicates downregulation.
