## Supplementary Figure 4 for "Deciphering Parasitic Strategies: Dual Transcriptomics Reveal Distinct Infection Mechanisms and Gall-like Traits in Rafflesiaceae"

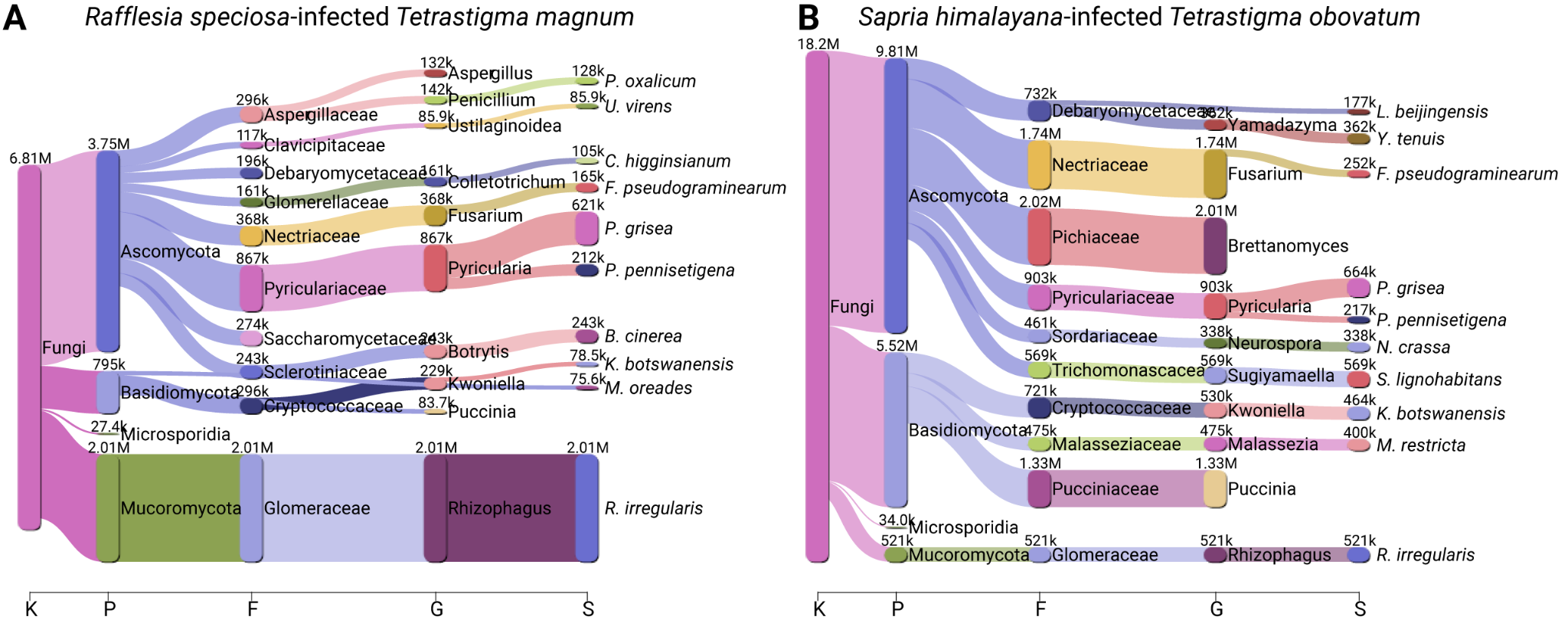

**Supplementary Figure 4: Taxonomic classification of fungal reads from interface tissue between Tetrastigma hosts and their parasites.** **A:** *Rafflesia speciosa*-infected *T. magnum*, **B:** *Sapria himalayana*-infected *T. obovatum*. Sankey diagrams show the flow of classified reads from kingdom (K) through phylum (P), family (F), genus (G), to species (S) level. Node width and connecting bands are proportional to read counts.
