## Supplementary Figure 5 for "Deciphering Parasitic Strategies: Dual Transcriptomics Reveal Distinct Infection Mechanisms and Gall-like Traits in Rafflesiaceae"

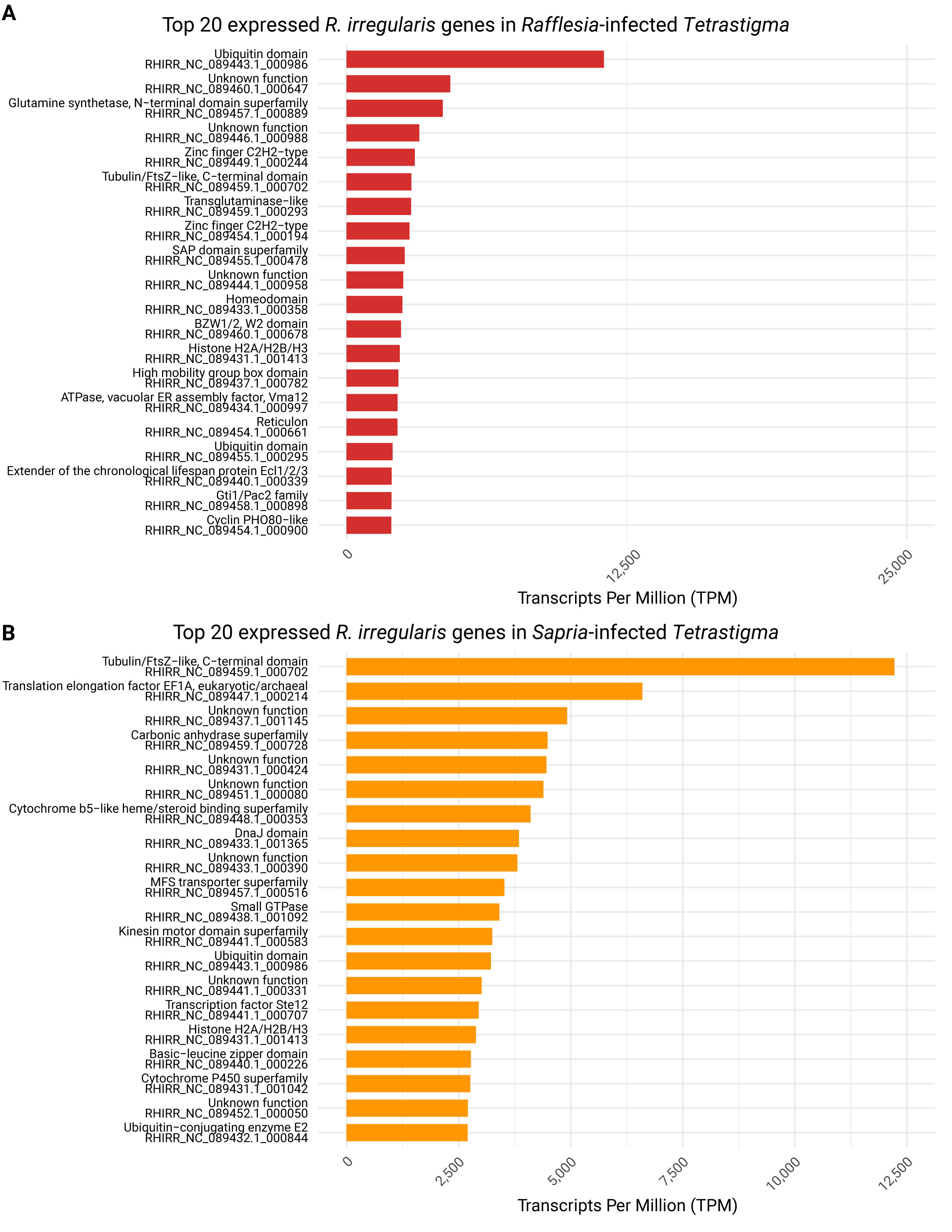

**Supplementary Figure 5: Top 20 *Rhizophagus irregularis* transcripts present in *Rafflesia*- and *Sapria*-infected *Tetrastigma* samples.**
