## Supplementary Figure 6 for "Deciphering Parasitic Strategies: Dual Transcriptomics Reveal Distinct Infection Mechanisms and Gall-like Traits in Rafflesiaceae"

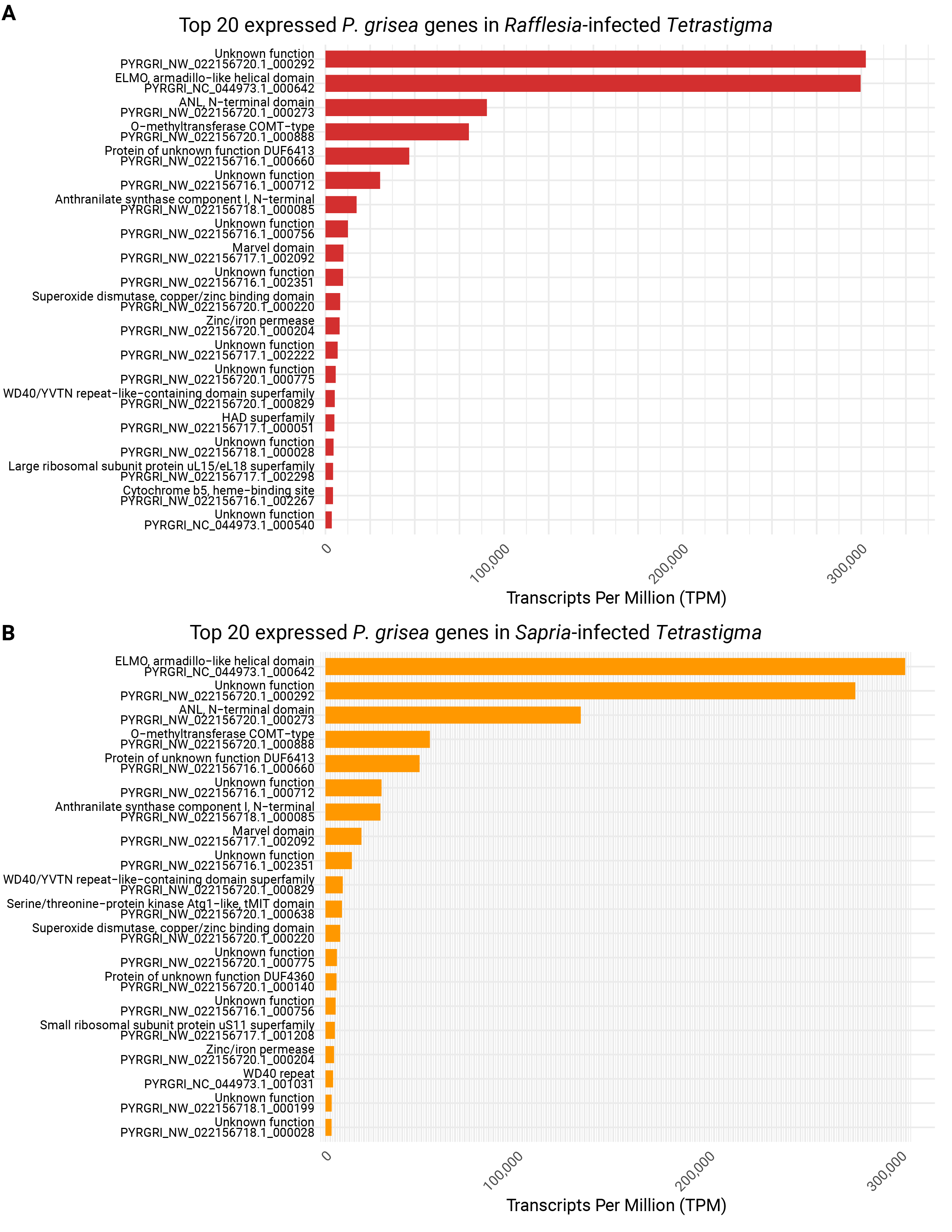

**Supplementary Figure 6: Top 20 *Pyricularia grisea* transcripts present in *Rafflesia*- and *Sapria*-infected *Tetrastigma* samples.**
