## Supplementary Figure 7 for "Deciphering Parasitic Strategies: Dual Transcriptomics Reveal Distinct Infection Mechanisms and Gall-like Traits in Rafflesiaceae"

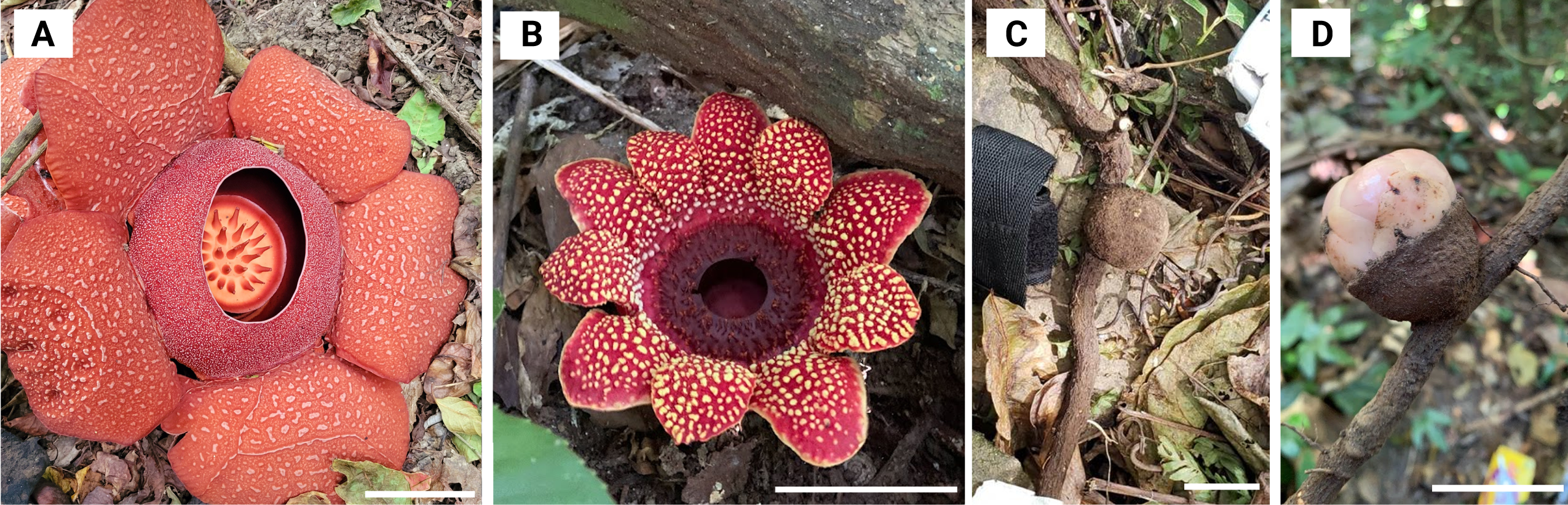

**Supplementary Figure 7: Plant samples used in this study.** Flower of **(A)** *R. speciosa* and **(B)** *S. himalayana*. The bud of **(C)** *R. speciosa* and **(D)** *S. himalayana* on *Tetrastigma* root. Scale bars are 10 cm for A, 5 cm for B, 2 cm for C, 2 cm for D.
